# Mapping the “missing” pathways of the anterior cingulate cortex in the human brain

**DOI:** 10.1101/2022.10.28.514260

**Authors:** Wei Tang, Javier Guaje, Shreyas Fadnavis, Eleftherios Garyfallidis

## Abstract

The anterior cingulate cortex (ACC) is functionally closely related with the insula and the ventral lateral prefrontal cortex (vlPFC). Extensive work on their functional relationships has led to the salience network theory and advanced understanding of value-based learning and decision making. However, the anatomical connections between the ACC and the two regions remain unknown in the human brain. Despite the anatomical ground truth established by nonhuman primate (NHP) tract-tracing, diffusion magnetic resonance imaging (dMRI) has not seen success identifying homologous pathways in humans. In this study we show that the negative finding does not reflect a cross-species discrepancy but rather a technical issue. We used NHP dMRI as a bridge to compare the ground-truth pathways in NHPs and dMRI-derived pathways in humans. The insight from NHP data helped pinpoint a bias in fiber orientation distribution functions (fODFs) caused by the disproportion of anterior-posterior vs. medial-lateral fibers in the human brain. Guided by this information, we successfully recovered the ACC-insula and ACC-vlPFC pathways that followed the same trajectories as in the NHP dMRI and tract-tracing data. Our findings provide an anatomical basis for the functional interactions among the ACC, the insula and the vlPFC.

## 1. Introduction

The anterior cingulate cortex (ACC) is a highly multifunctional region that plays an important role in cognitive and motivational control. Much of the insight on its functions is learned through regions that coactivate with the ACC in various cognitive tasks. Two of such regions, the insula and cytoarchitectonic area 47/12o in the ventrolateral prefrontal cortex (referred to as vlPFC here, see (Monosov & Rushworth, 2021) for review), share important roles with the ACC in value-based decision making (Hunt et al., 2018; Luk & Wallis, 2013), foraging behavior (Daw et al., 2006; Rushworth, Kolling, Sallet, Mars, 2012), attention reorientation (Dosenbach et al., 2006; Menon & Uddin, 2010) and pain perception (Critchley, 2005; Emmert et al., 2014; Rainville et al., 1997). The functional connectivity among the three regions indicates network interactions that support diverse behavior (Deen et al., 2011; Menon & Uddin, 2010; Taylor et al., 2009); the breakdown of such connectivity has been linked with psychiatric disorders (Szekely et al., 2017; Uddin, 2014; Whitton et al., 2019). However, despite the extensive functional studies, anatomical connectivity among the three regions in the human brain remains unclear. Studies have reported difficulty in finding white matter (WM) pathways between the ACC and the insula using tractography with diffusion magnetic resonance imaging (dMRI), the currently only noninvasive method for mapping structural connectivity *in vivo* (Cloutman et al., 2012). Likewise, reports on the ACC connections with the vlPFC have been scarce. Despite occasional mentioning of endpoint patterns (Ghaziri et al., 2017), the full fiber trajectory from the ACC to the vlPFC remains unknown.

The unresolved anatomical puzzle posts challenges in interpreting the functional findings and hinders clinical applications. The well-known salience network theory builds upon the hypothesis that the ACC closely interacts with the insula (Menon, 2015; Menon & Uddin, 2010; Seeley, 2019) to detect salient stimuli and switch between exogenous and self-related information processing. The lack of anatomical connections has led to a hypothesis that there is a third region (or set of regions) mediating the observed functional interactions (Cloutman et al., 2012). Moreover, in clinical interventions for treatment-resistant psychiatric disorders, such as deep-brain stimulation, the precise location of fiber connections is crucial for identifying stimulation targets (Sullivan et al., 2021). Thus, the negative finding of ACC-insula and ACC-vlPFC pathways not only challenges the understanding but also practices of related network theories. Nevertheless, despite the negative finding in humans, tract-tracing studies in nonhuman primates (NHPs) have long established strong bidirectional projections among the three regions (Carmichael & Price, 1996; Schmahmann & Pandya, 2006; Trambaiolli et al., 2022; Vogt & Pandya, 1987). Schmahmann and Pandya (2006) detailed the pathways of ACC that traverse laterally through the deep WM in the frontal lobe to reach the vlPFC and through the extreme capsule (EmC) to reach the insula. These fiber trajectories are replicable by dMRI tractography in NHPs *in vivo*: streamlines derived from ACC and vlPFC seeds have shown mutual connections to the other region (Neubert et al., 2015), and the dMRI-derived EmC has shown terminations in the ACC (Mars et al., 2016), both consistent with the tract-tracing work. These results, however, have not seen replication in humans (in Neubert et al., 2015, for example, the ACC-vlPFC connectivity was rather dissimilar between the human and macaque results even though the overall connectivity fingerprints were similar).

It is yet too early to conclude that the pathways found in NHPs do not exist in humans. Recent work has reported encouraging success in finding cross-species homology for major fiber bundles, while pointing to technical issues that can cause tractography to dismiss anatomical ground truth (Girard et al., 2020; Grier et al., 2020; Maffei et al., 2021). Indeed, there is a biological basis for possible tractography failure in the ACC-related pathways. The deep WM region that encompasses the ACC-vlPFC and ACC-insula fibers is occupied by thick anterior-posterior (A-P) running bundles. These bundles take up a substantial portion of the WM volume and dominate its dMRI signals. Thus, fiber orientation models tend to assign disproportionally high probability towards the A-P direction and low probability towards the medial-lateral direction, causing erroneously low sampling rate for catching the medial-lateral fibers. This problem can be more prominent for human tractography because the problematic WM region in the human frontal lobe is much enlarged compared to NHPs’.

In this study, we examine and tackle the problem that causes the “missing” ACC pathways in humans. We used NHP dMRI as a bridge to compare the ground-truth pathways from NHP tract-tracing data and dMRI-derived pathways in the human brain. We show that while dMRI tractography in NHPs faithfully replicated the tract-tracing data, the same standard tractography method could not capture the same pathways in humans. Informed by the NHP dMRI data, we identified a technical issue that impairs the estimation of fiber orientation distribution functions (fODFs) in the human brain. We developed a fix to the problem based on the assessment of signal-to-noise ratio (SNR), and recovered the ACC-vlPFC and ACC-insula pathways that followed the same trajectories as in the NHP dMRI and tract-tracing data. Our findings demonstrate the effectiveness of a cross-species, multimodal approach to delineate detailed fiber trajectories in the living human brain, opening the door to validate the anatomical basis underlying known functional networks.

## 2. Methods

### 2.1. Participants and procedure

T1-weighted images and dMRI data analyzed in this study were freely obtained from the Human Connectome Project (HCP) via ConnectomeDB (https://db.humanconnectome.org) and Primate Data Exchange (PRIME-DE) (Milham et al., 2018). The details of participants, procedure and MRI acquisition has been documented in corresponding publications (Folloni et al., 2019; Glasser et al., 2013). Briefly, a total of 174 individuals who had completed diffusion and structural scans were included from the subject pool included in the HCP 7 T dataset. No participants were excluded. There are 67 males and 107 females in the 174 participants, with ages mainly in the range 22–36 years. The NHP dMRI data contained post-morten scans of 6 macaque monkeys (*Macaca Mulatta*, age 4.03-15.81 years, mean age 9.98, standard deviation 4.64, two females). The post-mortem tissue preservation procedure has been documented on PRIME-DE (https://fcon_1000.projects.nitrc.org/indi/PRIME/oxford2.html) and in (Folloni et al., 2019). Briefly, immediately after death, the brains were perfusion fixed with formalin. Approximately one week before MRI scanning, the brains were perfused in phosphate buffer solution to enhance their diffusion signal. During scanning, the brains were suspended in fomblin.

### 2.2. MRI acquisition and preprocessing

Details of the human 7 T diffusion and T1w image acquisition protocols are provided in the HCP reference manual (https://humanconnectome.org/study/hcp-young-adult/document/1200-subjects-data-release). Briefly, the dMRI scans used a monopolar scheme with single-shot 2D spin-echo multiband (factor=2) EPI acquisition, with the main parameters as follows: 1.05mm isotropic voxel (FOV = 210 × 210 mm^2^, matrix size=200 × 200), 132 transversal slices acquired in interleaved order without a gap, phase encoding applied along the anterior–posterior direction, phase encoding acceleration (GRAPPA) factor 3, two shells with b-values=1000, 2000 s/mm^2^, repetition time/echo time=7000/71ms, 65 unique diffusion gradient directions and 6 b0 images were obtained for each phase encoding direction pair (AP and PA pairs). The total scanning time for the dMRI protocol was about 40 min. The preprocessing was performed as described by (Glasser et al., 2013), based on the updated diffusion pipeline (v3.19.0), including basic preprocessing, distortion correction, eddy current correction, motion correction, gradient nonlinearity correction, and registration of the mean b0 image to native T1w images with FLIRT BBR+bbregister and transformation of dMRI.

The NHP dMRI acquisition and preprocessing pertained to the public dataset are documented on PRIME-DE (https://fcon_1000.projects.nitrc.org/indi/PRIME/oxford2.html) and in (Folloni et al., 2019). Briefly, diffusion-weighted imaging was done on a 7T superconductive magnet driven by an Agilent DirectDrive console (Agilent Technologies, Santa Clara, CA, USA) using a 72 mm ID quadrature birdcage RF coil (Rapid Biomedical GmbH, Rimpar, Germany). Diffusion-weighted images were acquired by a spin-echo single line readout protocol (DW-SEMS: TE=25 ms; TR=10 s; matrix size=128 × 128; resolution=0.6 × 0.6 mm; 128 axial slices; slice thickness=0.6 mm). Each subject’s dataset consisted of 16 non-diffusion-weighted (b = 0 s/mm^2^) and 128 diffusion-weighted (b = 4000 s/mm^2^) volumes with diffusion directions homogeneously distributed over the sphere. For preprocessing, eddy current correction was applied using the FDT tool by FSL.

### 2.3. Fiber orientation modeling and tractography

For both the human and NHP data, the same fiber orientation model and tractography methods were applied. All the analyses were implemented using DIPY (Garyfallidis et al., 2014). To estimate the fODF, a constant solid angle (CSA) model (Aganj et al., 2010) was fitted to each voxel with a spherical harmonic order of 6, resulting in a density function of 362 directions. Anatomically constrained tractography (ACT) (Smith et al., 2012) was applied, which uses tissue priors for the gray matter (GM), WM and cerebrospinal fluid (CSF) to determine the streamline trajectories and termination points during tracking. Streamlines were generated from the seed masks with a seed density of 6 and tracked until they reached the GM in either the cortex or subcortical structures. Streamlines were discarded if they entered CSF or showed an excessive curving angle (< 30º) in the WM. A probabilistic method based on particle filtering (Girard et al., 2014) was used to generate the streamlines. To facilitate visualization, the 3D streamlines in each individual brain in all the figures were generated using deterministic tracking (Garyfallidis et al., 2014). While deterministic tracking was used for visualization, all the streamline statistics were calculated based on the probabilistic-tracking results.

### 2.4. Region masks for tracking and assessing streamlines

A seed mask in the ACC was created for generating streamlines and two regions of interest (ROIs) in the WM were used for assessing the streamline trajectories. All three masks were first created in a template brain (for NHP, the NMT template (Jung et al., 2021); for humans, the MNI152 1 mm T1 brain (Fonov et al., 2009)), and then registered to each subject’s native space. Cross-species homology was a priority in determining the mask locations. The seed mask encompassed the most anterior region of the ACC. In NHPs, the center of the mask was the rostral end of the cingulate sulcus. In humans, area 32 in the Petrides atlas (available as supplemental in (Tang et al., 2019)) was divided into 10 equal-volume subregions. Starting from the most superior subregion, the seventh subregion was used as the seed mask.

To assess and compare streamline trajectories within and between species, two WM masks were created, separately for each species. The first mask covered the deep WM region in the frontal lobe (ROI 1) between the ACC and the vlPFC; the second mask encompassed a segment of the EmC (ROI 2) adjacent to the insula. Both masks were defined based on six anatomical landmarks demarcating the anterior, posterior, superior, inferior, left and right boundaries. For ROI 1 landmarks, anterior: the most anterior slice of area 24; posterior: the most anterior slice of the striatum; superior: the most superior slice of the principal sulcus in NHPs and that of the inferior frontal sulcus in humans; inferior: the cortical GM/WM boundary on each coronal slice; left: the leftmost slice of area 13; right: the cortical GM/WM boundary on each coronal slice. For ROI 2 landmarks, anterior: the most anterior slice of the insula; posterior: the most anterior slice of the amygdala; superior: the most superior slice of the insula; inferior: the most inferior slice of the insula; left: the striatal GM/WM boundary on each slice; right: the insular GM/WM boundary on each slice.

### 2.5. Fiber orientation angle analysis

To compare the estimated probability of A-P oriented and medial-laterally oriented fibers in a chosen voxel, we define the orientation angle *ϕ* as follows: For a vector *d* in a given fODF, define its projection in the axial plane (spanned by the A-P and right-left (R-L) axes) as *d*′. Angle *ϕ* is the angle between *d*′ and the R-L axis (Fig. 1A). Its value range is defined as follows (Fig. 1B): *ϕ* equals 0 when *d*′ completely aligns with the A-P axis regardless whether *d*′ points to the A- or P-direction; *ϕ* takes a negative sign and decreases from 0 to -90 as *d*′ move towards the R-half of the R-L axis; *ϕ* equals -90 when *d*′ completely aligns with the R-half of the R-L axis. Correspondingly, *ϕ* takes a positive sign and increases from 0 to 90 when *d*′ moves towards the L-half of the R-L axis; *ϕ* equals 90 when *d*′ completely aligns with the L-half of the R-L axis. Because the value is symmetric for the A- and P-half of the A-P axis, *ϕ* ranges in the interval [-90, 90].

**Figure 1.**
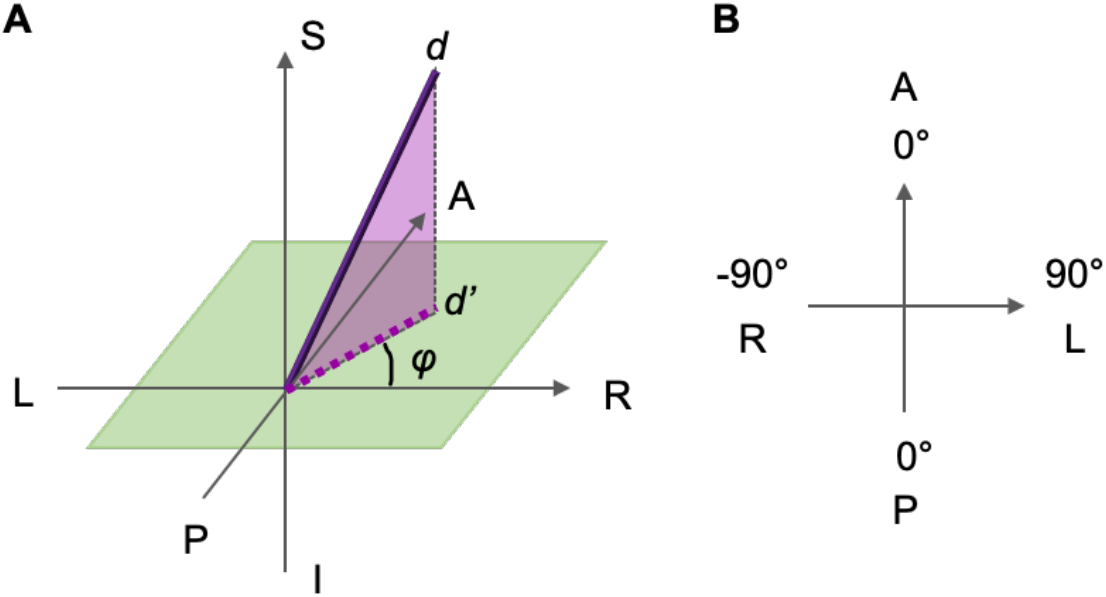
Definition of the fiber orientation angle. (A) Three-dimensional view of the orientation vector *d* and its associated angle *ϕ*. (B) In-plane view of *ϕ* and its value range.

### 2.6. Cross-subject variance weighting

For each vector *d* of the fODF from each voxel in the ROI 1 mask, we computed the cross-subject standard deviation of the associated probability values. Then, in each subject, the probability value associated with each *d* was divided by its corresponding cross-subject standard deviation.

### 2.7. Orientation angle histogram

To assess how the SNR-based rescaling affect fODFs in different directions, histograms of *ϕ* were computed as follows: (1) In each subject, for each voxel in ROI 1, the probability values of the original fODF was subtracted from the probability values of the rescaled fODF in each direction. (2) The fODF differences were categorized into increase (positive difference) and decrease (negative difference). (3) In each subject, separately for the increased and decreased fODFs, the difference values were pooled over the 362 directions and all the voxels in ROI 1. The median of each pooled population was calculated for threshold. (4) In each subject, for increased fODFs, the vector *d*s associated with above-median differences were identified. A histogram of *ϕ* angles associated with these vectors was computed over 20 equal-sized bins spanning the [-90, 90] interval. For decreased fODFs, the histogram was calculated in the same manner but for vector *d*s associated with below-median values. (5) The histograms were then averaged across subjects.

### 2.8 Streamline density and surface projection

Streamline density in each voxel was calculated as the number of streamlines passing that voxel divided by the total number of valid streamlines generated. For visualization on the FreeSurfer-generated cortical surfaces, in each subject, the density scores between the pial surface and up to 2 mm into the WM were projected to the surface space using the FreeSurfer mri_vol2surf utility. Individual cortical maps were morphed to the fsaverage space for computing and visualizing the group average map.

## 3. Results

### 3.1. Anatomical pathways in NHPs

The fiber pathways connecting the ACC with the vlPFC and the insula have been identified using tract tracing in NHPs (Carmichael & Price, 1996; Schmahmann & Pandya, 2006; Trambaiolli et al., 2022; Vogt & Pandya, 1987). According to previous studies, fibers from the ACC take a medial-to-lateral trajectory that traverses the deep WM to reach the vlPFC (Fig. 2A, top). Fibers that target the insula take a medial-to-lateral path to join the EmC (Fig. 2A, bottom), which then travel posteriorly to reach the insula. To test whether diffusion tractography can capture these known pathways in the same species, we placed seeds in matching positions to that of the ACC injection shown in Fig. 2A and examined the spatial distribution of streamline density generated from the NHP dMRI data (Fig. 2B). Consistently, we observed that the tractography-generated streamlines traversed the same WM regions to connect with the vlPFC and the insula (Fig 2B). In the 3D view of streamline trajectories (Fig. 2C), the streamlines traveled laterally to reach the vlPFC region in the same coronal plane (Fig. 2C, middle). A separate group of streamlines first traveled laterally and then caudally to join the EmC, providing a bundle to reach the insula and posterior regions (Fig. 2C, bottom).

**Figure 2.**
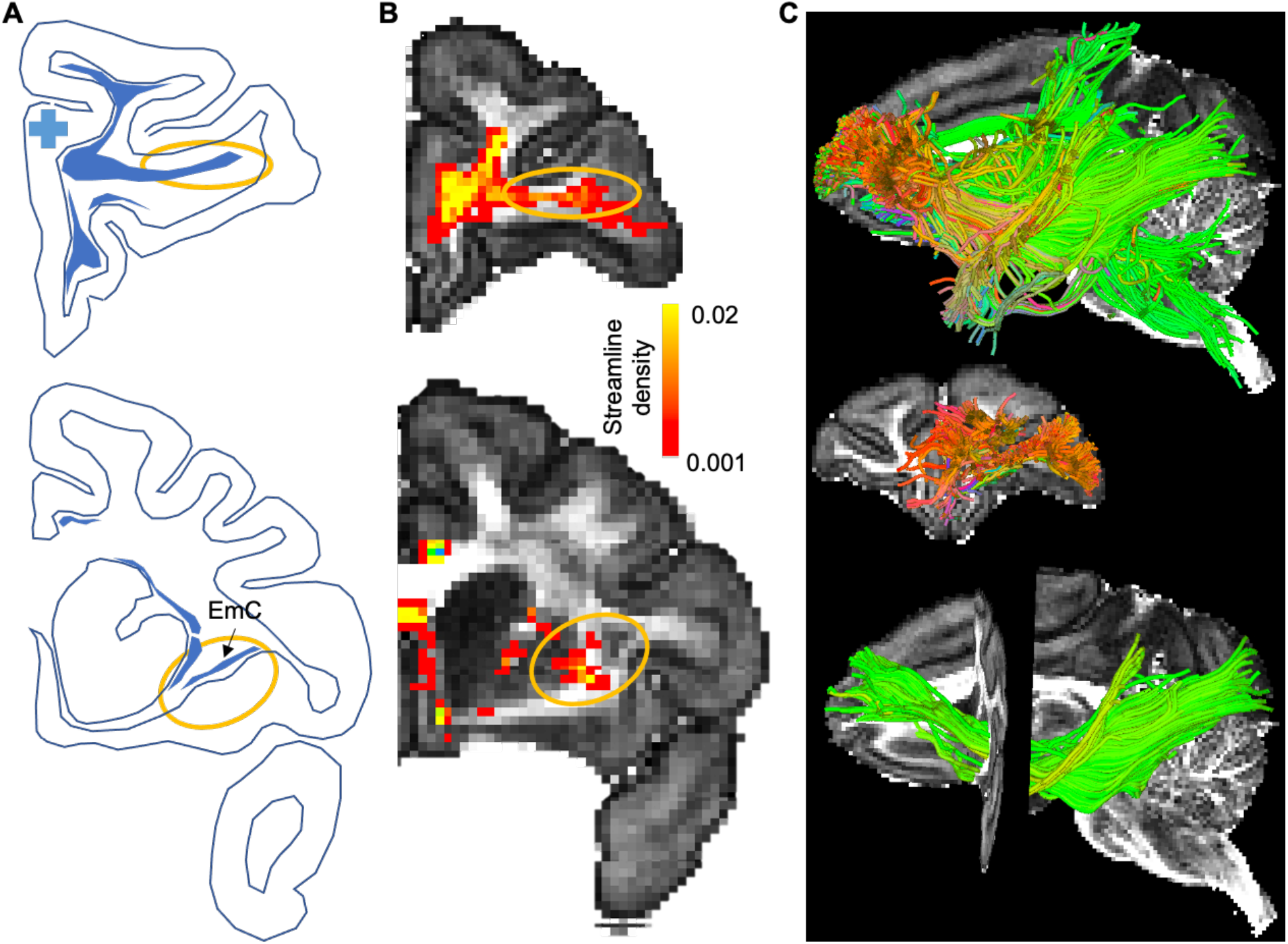
Tractography-derived fiber bundles in comparison with the anatomical ground truth. (A) Fiber trajectory masks created following Schmahmann & Pandya (2006), Case 30, showing two exemplar coronal slices. Blue cross indicates the tracer injection site in rostral area 24. Filled blue fields indicate fiber trajectories labeled by the tracer. Orange ellipses highlight the pathways that the ACC fibers use to reach the vlPFC and the insula. (B) Tractography-derived streamlines from a seed at the location matching the injection site in A, shown on two coronal MRI slices with matching z-coordinates of the slices in A. Colors represent the streamline density. Orange ellipses highlight the pathways into the vlPFC, consistent with the ground truth identified by tract-tracing. (C) A 3D visualization showing all the streamlines in a sagittal view (top), the streamlines reaching the vlPFC in a coronal view (middle) and the streamlines of the EmC in a sagittal view (bottom).

### 3.2. Absence of the ACC pathways in human tractography

Applying the same tractography method to the HCP data did not result in homologous pathways in the human brain. We placed a seed in the human cingulate area 24, a location that matches the seed in the NHP data (Fig. 3A). We generated streamlines using the same fODF modeling and tracking methods. In contrast to the results in the monkey, very few streamlines reached the vlPFC (Fig. 3B & C). The streamline density was low to none in the WM near the vlPFC or in the EmC (Fig. 3B). Indeed, the entire EmC was missing in 122 of the 174 HCP subjects (Supplemental Figures). To compare the tractography results across species, we calculated the streamline density in two WM ROIs (Fig. 3D), one situated between the cingulate gyrus and the lateral prefrontal cortex (ROI 1), the other between the putamen and the insula cortex (ROI 2). The locations of the ROIs were determined by landmarks chosen to maximize cross-species homology (see Methods). To control for systematic differences of streamline density across species, we divided the cross-voxel average density in each ROI by the cross-voxel average density in the entire WM in each species. In both ROIs, the normalized density showed highly significant differences across species (Fig. 3E). The density in the NHP (averaged across 7 animals) was significantly higher than the density in humans (averaged across 174 individuals) (independent-sample *t* tests, ROI 1: *t* = 36.70, *p* < 1× 10^−10^; ROI 2: *t* = 16.88, *p* < 1× 10^−10^).

**Figure 3.**
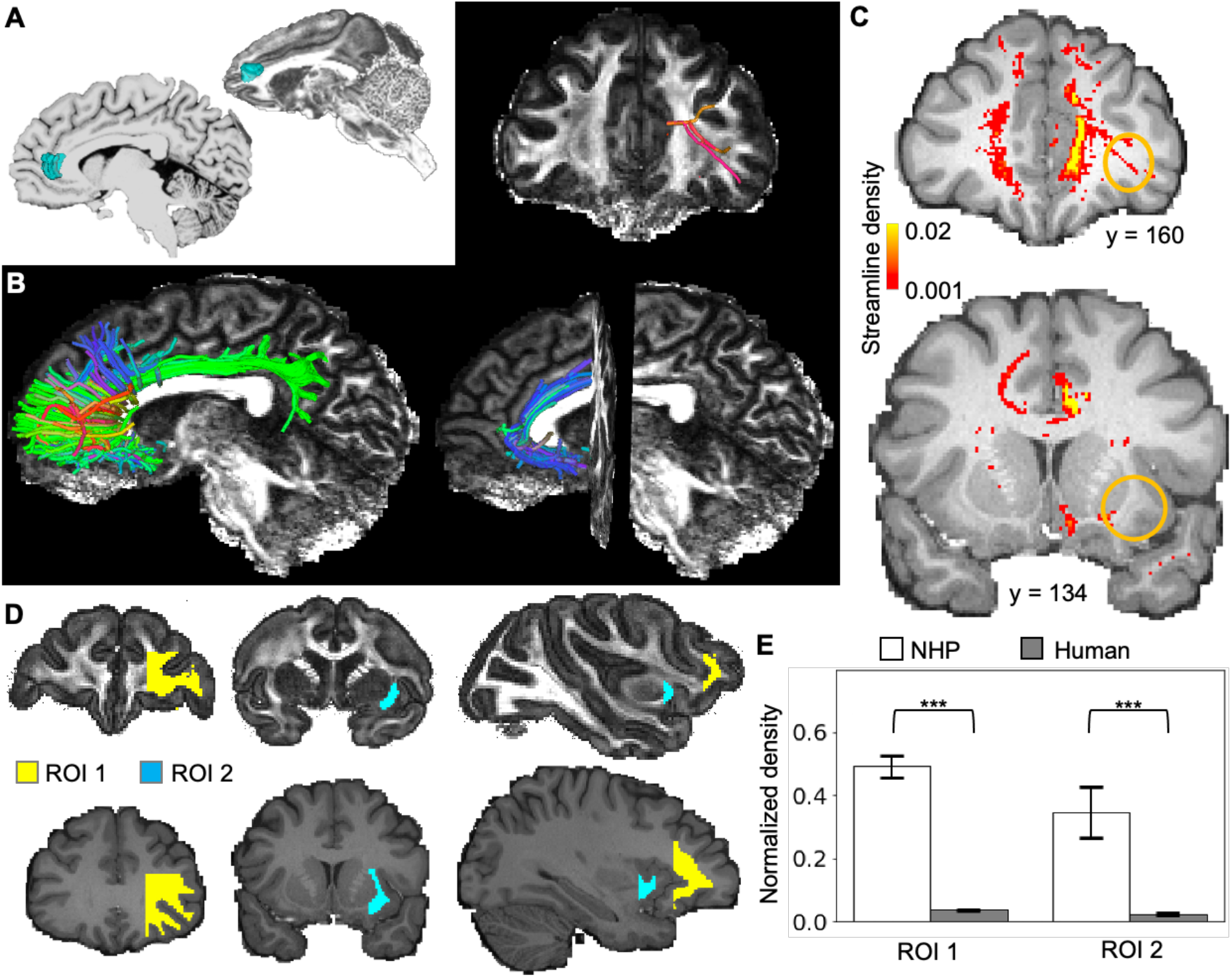
Absence of medial-lateral connections in human dMRI tractography. (A) ACC seed locations in the human (left) and macaque (right) brain, shown in blue in an individual subject of each species. (B) Tractography result from a single HCP subject (100610), showing all the streamlines in a sagittal view (bottom left), the streamlines reaching the vlPFC in a coronal view (upper right) and the streamlines traveling ventrally in a sagittal view (bottom right). (C) Streamline density on two representative slices showing the low-to-none density in the deep WM and EmC marked by orange circles. (D) Locations of ROIs 1 & 2 in the NHPs and humans. (E) Normalized streamline density (log-transformed for statistical comparison) of ROIs 1 & 2 in the NHPs and humans. ***: *p* < 1× 10^−10^.

### 3.3. Structural differences in the deep WM between the human and NHP brains

To identify potential causes of the low-to-none streamline density in the human subjects, we examined the fODFs in the deep WM of ROI 1. Visual inspection suggested that the fODFs in humans had a strong bias towards the A-P axis, in contrast to the more uniform fODFs in NHPs. To illustrate, we extracted fODFs from a coronal slice in a macaque and a human brain (Fig. 4A middle panel, MR images), with the voxel locations matched across species. Compared with the fODFs in the macaque (Fig. 4A left panel), fODFs in the human (right panel) showed high probability mass along the A-P axis and much lower probability mass in the medial-lateral direction. To examine whether this difference was systematic across species, for each subject of each species, we computed the histogram of orientation angles (see Methods for the definition of *ϕ*) associated with above-median probability in the fODFs. The angle *ϕ* of voxels in ROI 1 were pooled for computing each histogram, and the histograms were averaged across subjects in each species. In NHPs, the orientation angles associated with above-median probability was evenly distributed over the interval [− 90°, 90°], while in humans high counts tended to center around (Fig. 4A middle panel, bottom plot). This pattern suggested that high probability in the A-P direction (*ϕ* = 0°) was more frequently observed in humans than in NHPs. The A-P bias in humans potentially explains why the medial-lateral streamlines found by NHP tractography were not replicated in humans.

**Figure 4.**
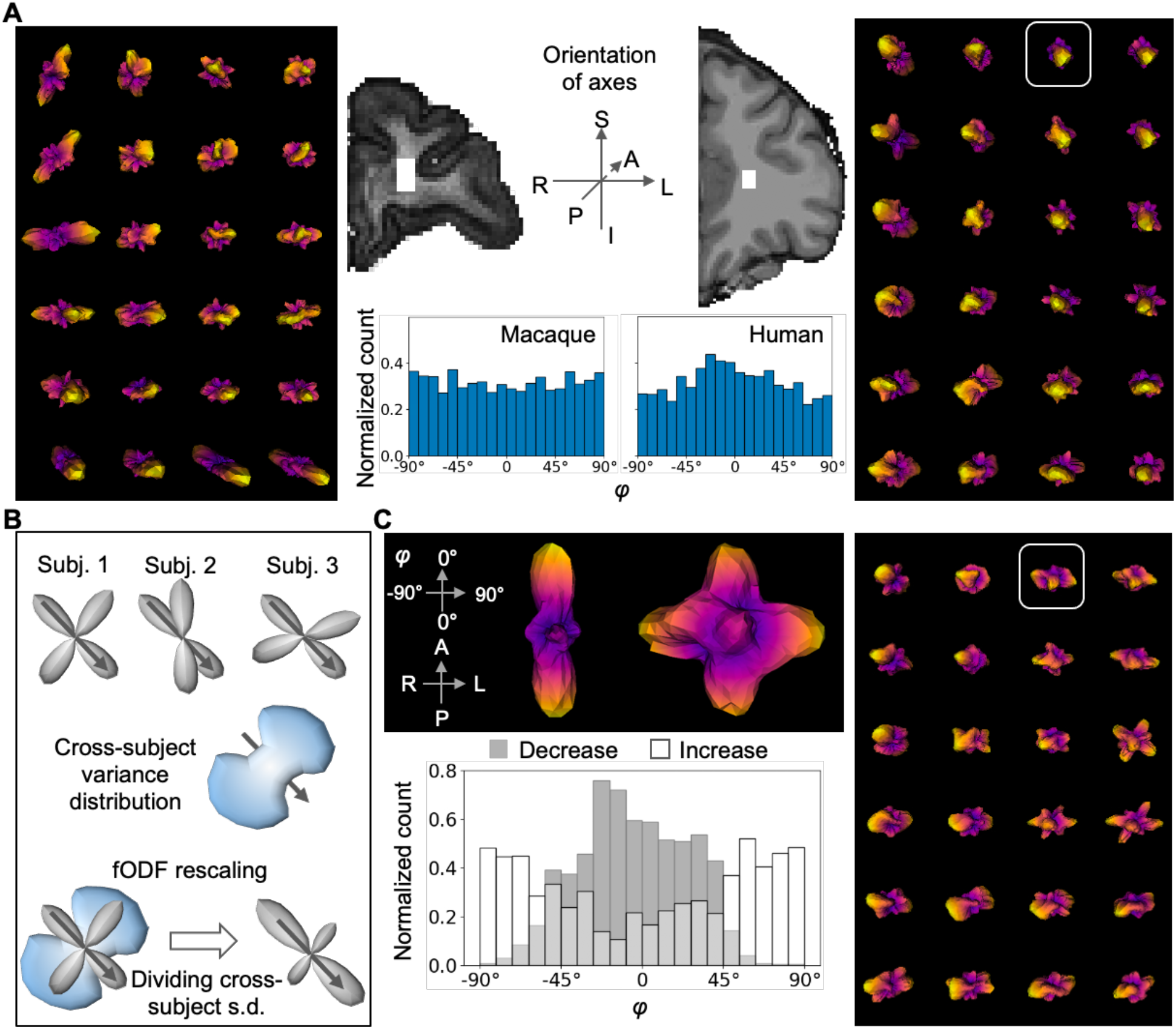
Rescaling fODFs to boost the probability in the directions of existing fibers. (A) Comparison of fODFs in the NHP and human brains. Middle panel shows two coronal slices from an NHP and a human brain, respectively, with the white masks marking the voxels for comparison. The fODFs of voxels from each mask are shown in the left and right panels, respectively. The axes of the 3D fODF display are shown by the schematic in the middle panel. The histograms of angle is shown at the bottom. Abbreviations: A=anterior, P=posterior, S=superior, I=inferior, L=left, R=Right. (B) Schematics showing the rationale of SNR-based rescaling of fODFs. Top: Schematic example of the variance across three fODFs from three subjects. Gray arrow marks the density resulting from the dMRI signal of a true fiber bundle running in the pointed direction, and this bundle consistently exists in each subject. Abbreviation: Subj.=subject. Middle: Schematic variance function across the three fODFs. The variance is low in the arrow-pointed direction because presumably the true fiber bundle generates consistent dMRI signals across subjects. The variance in the other directions is high because presumably the density in these directions are resulted from random noise. Bottom: Schematic of the fODF rescaling. Density in the arrow direction proportionally increases after *ϕ* the fODF is divided by the cross-subject s.d. (C) Rescaling applied to the fODFs of the same example human subject in A. Rescaled fODFs from the same voxels are shown in the right panel. Middle top: the rescaling effect on a single fODF from the voxel highlighted in the right panels (left: before rescaling, right: after rescaling). The fODF is reoriented for visualizing the density in the R-L direction. Middle bottom: Histogram of *ϕ* associated with increased and decreased probability due to rescaling

To our assessment, the fODF A-P bias was a resolvable technical issue rather than reflecting the biological truth of fiber orientations in ROI 1. The WM volume is much enlarged in humans compared to NHPs, and the A-P oriented bundles are increasingly disproportionate in size compared to the medial-laterally oriented fibers. Thus, the lack of medial-lateral streamlines did not necessarily indicate a lack of medial-lateral fibers but possibly the difference in SNR between the A-P and medial-lateral directions. The tracking algorithm might have sampled frequently in the A-P direction while failing to distinguish fibers from noise in the medial-lateral direction. In order to address this issue, we aimed to distinguish fODF peaks generated by axons from those generated by random water diffusivity. Our strategy was to rescale the fODF by an SNR-based metric. Theoretically, if in every subject there were axons passing through the same brain region in the same direction, then the fODF peaks in this region and direction would be relatively consistent across subjects, while peaks driven by noise would randomly vary (Fig. 4B, top). Thus, a low cross-subject variance would reflect high SNR, meaning that the dMRI signal in the corresponding direction was likely driven by existing axons (Fig. 4B, middle). Following this rationale, in each subject and for each voxel in ROI 1, we divided the fODF by the cross-subject standard deviation (s.d.) (Fig. 4B, bottom). If the medial-lateral pathways indeed exist in every subject, the probability in the direction of existing fibers would be scaled up and those of random water diffusivity would be scaled down (Fig. 4B, bottom).

Validating our speculation, the rescaling resulted in increased probability in the medial-lateral direction (Fig. 4C). To illustrate this effect, we plotted an example fODF from a single voxel (Fig. 4C, middle panel). Before rescaling (left), the fODF peaks were highly aligned with the A-P axis; after rescaling (right), additional peaks were observed along the orthogonal R-L axis. Similar effects can be seen in most of the voxels shown in the right panel of Fig. 4C: after rescaling, many fODF peaks became visible in the plane orthogonal to the A-P axis. To test whether this was a general effect across voxels and subjects, we measured the histogram of orientation angles in which the fODF showed above-median increase or decrease after rescaling (see Methods). The decrease was mostly found around 0° (A-P axis) while the increase was mostly found around ± 90º (R-L axis) (Fig. 4C, middle panel).

### 3.4. Tractography with rescaled fODFs

After rescaling the fODFs of ROI 1 in each subject’s native space, we regenerated streamlines from the same ACC seed using the same tractography method. We first compared results in the same example subject before and after fODF rescaling (Fig. 5). Visual inspection suggested a fundamental change in the streamlines connecting the seed and the vlPFC and streamlines in the EmC (Fig. 5A). After rescaling, a full bundle oriented in the medial-lateral direction was observed to connect the seed and the vlPFC (Fig. 5A, upper right). Moreover, the EmC was clearly identifiable (Fig. 5A, lower right). The before-after effect was also prominent in the density map (Fig. 5B). For both ROIs 1 & 2, the increase of average streamline density across voxels was highly significant (Fig. 5C) (independent-sample *t* tests, ROI 1: *t* = 65.22, *p* < 1× 10^−10^; ROI 2: *t* = 16.00, *p* < 1× 10^−10^). Next, we examined the before-after effects in all of the 174 HCP subjects. At the group level, consistent with the example-subject result, the increase of average streamline density across subjects was highly significant for both ROIs 1 & 2 (Fig. 6A) (independent-sample *t* tests, ROI 1: *t* = -24.25, *p* < 1× 10^−10^; ROI 2: *t* = -17.61, *p* < 1× 10^−10^). Visualization of the subject-average density on the cortical surface verified that the increase of streamlines was in the vicinity of the vlPFC and the insula (Fig. 6B). At the individual level, we examined the recovery of EmC in each of the 122 subjects for whom the EmC was not found before fODF rescaling. This effect is illustrated in Fig. 7, in which we plotted the full tractograms of 10 representative subjects before and after fODF rescaling. In each subject, the EmC went from nonexistent to a full-body recovery. Indeed, the EmC was recovered in 115 out of the 122 subjects (Supplemental Figures).

**Figure 5.**
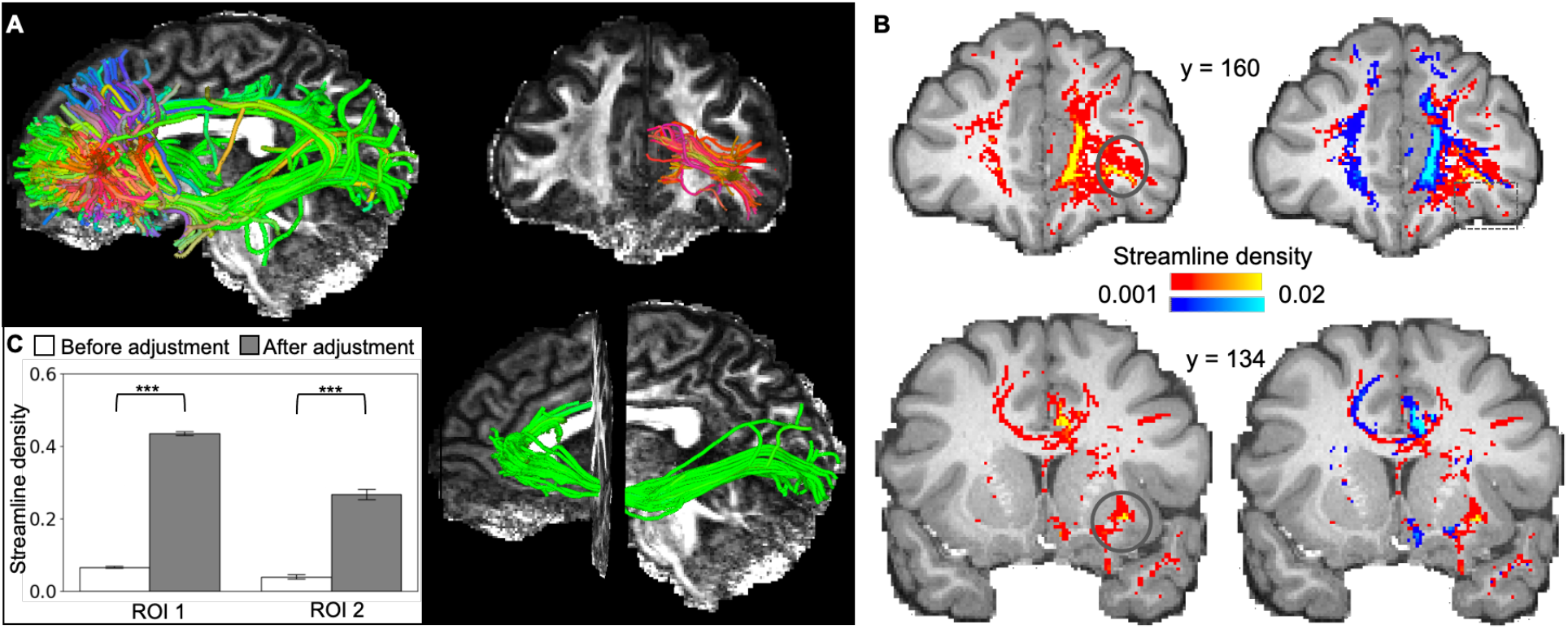
Tractogram after fODF rescaling in an example human subject. (A) Streamlines generated by tractography in 3D visualization, showing a global view (top) and the streamlines reaching the vlPFC in coronal (middle) and sagittal (bottom) views. (B) Tractography-derived streamline trajectories from the ACC seed, shown on two coronal MRI slices with matching z-coordinates of the slices shown in A. Colors represent streamline density. Orange ellipses highlight the pathways into the vlPFC, consistent with the ground truth identified by tract-tracing. (C) Streamline density (log-transformed) averaged across voxels in ROIs 1 & 2. Error bar represents standard error across voxels. ***: *p* <1× 10^−10^.

**Figure 6.**
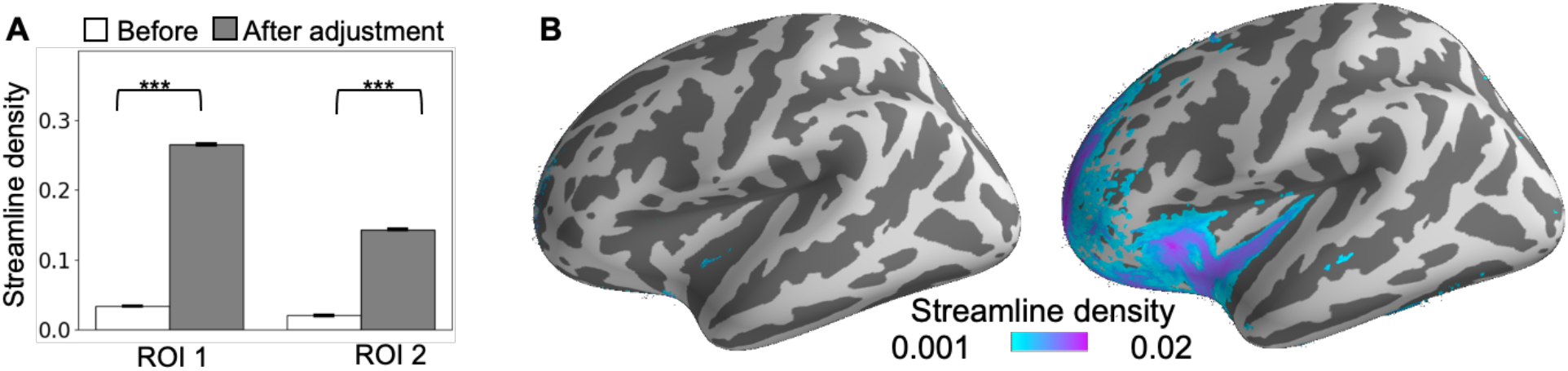
Cross-subject average of streamline density before and after fODF rescaling. (A) Mean streamline density (log-transformed) across subjects in ROIs 1 & 2. ***: *p* <1× 10^−10^. (B) Streamline density projected on the cortical surface before (left) and after (right) fODF rescaling.

**Figure 7.**
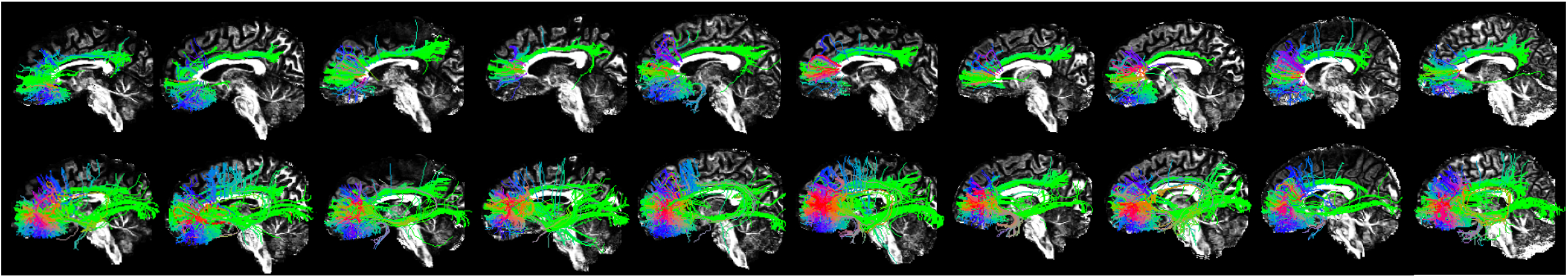
Tractograms before (top) and after (bottom) fODF rescaling in 10 subjects.

## 4. Discussion

Identifying pathways connected with the cingulate cortex has been a challenge for dMRI tractography. The cingulum bundle and other A-P oriented large bundles tend to cause predominantly high probability mass in the fODFs. As a result, streamlines connecting the medial wall with the lateral cortex are difficult to track and prone to false negative outcomes. This issue can lead to erroneous understanding of anatomical connections and related functional interpretations of the ACC and other brain regions. In this report, we show that the “missing” connections between the ACC and the vlPFC and the insula are indeed a false negative that can be corrected by rescaling fODFs based on the SNR estimation. By consulting the ground truth in NHP tract-tracing data, and using NHP tractography as an intermediate step to validate the tractography methodology, we were able to draw intuition of the biological cause for tractography to fail in humans. The recovery of the ACC-vlPFC and ACC-insula pathways demonstrates the promise of a cross-species, multimodal approach in precisely mapping anatomical connections in the human brain.

### 4.1 Anatomical connectivity underlying the salience network

Our findings open the door to exploring the anatomical basis of the salience network. The classical definition of the salience network (Menon & Uddin, 2010) highlights two core nodes, the dorsal ACC and the insula. While their functional relationships have been extensively discussed in the literature (Uddin, 2014), the anatomical connections to support their interactions remain unclear in humans (Cloutman et al., 2012). A potential cause of this problem is the difficulty to track streamlines between the medial and lateral cortices as described above. Here we show that the EmC, a bundle running longitudinally and adjacent to the insula, is joined by fibers from the ACC. The ACC seed was placed rostral to the classical dorsal ACC node of the salience network, due to homology considerations for cross-species comparisons. Nonetheless, the success in tracking streamlines from the rostral ACC seed is straightforwardly applicable to dorsal ACC regions. Fibers originating from the frontal medial wall share the same deep WM volume (i.e. ROI 1) to reach the lateral cortex. Thus, the fODF rescaling in ROI 1 can be a common fix for tracking streamlines that connect different ACC subregions and the insula, including nodes of the salience network.

Another relevant finding is the connections between the ACC and the vlPFC. While the majority of salience network studies are focused on the ACC and insula, whether the vlPFC is also a node or not remains an unresolved question (Trambaiolli et al., 2022). In value-based decision making, the functions of the vlPFC and dorsal ACC are closely related in both humans and NHPs (Daw et al., 2006; Hunt et al., 2018; Luk & Wallis, 2013; Monosov & Rushworth, 2021). Whether the dorsal ACC and possibly the vlPFC in these studies are part of the salience network will bring fundamental insights to advance both fields. This question cannot be resolved with functional data alone because functional interactions can arise from indirect anatomical connections. In the NHP tract-tracing literature, strong bidirectional projections have been established among the ACC, vlPFC and insula (Carmichael & Price, 1996; Schmahmann & Pandya, 2006; Trambaiolli et al., 2022; Vogt & Pandya, 1987). Here we show that some of these anatomical pathways can be replicated in humans using dMRI tractography. Importantly, we found both vlPFC and insula connections with the same ACC seed. The approach can be directly applied to the other ACC subregions to explore whether vlPFC and insula connections converge onto an ACC subregion that is part of the salience network.

### 4.2 Anterior origin of the EmC in the medial PFC

Previous work on dissecting the EmC with dMRI tractography have been focused on its temporal and occipital segments (e.g. (Makris & Pandya, 2009; Mars et al., 2016; Radwan et al., 2022)), and not much discussion was on terminations in the frontal cortex. Our results suggest that fibers from the ACC also join the EmC in the human brain, adding an important criterion for segmenting this bundle. The EmC is the major pathway for the insula and superior temporal regions to connect with the rest of the cortex. Identifying the precise fiber composition in the EmC, including the regions they connect with, their spatial topography within the bundle, and their trajectory traversing the WM, is essential for understanding the EmC-connected functional networks and related brain disorders. Adding ACC fibers to the EmC calls for a re-consideration of its segmentation in future work.

### 4.3 SNR measurements for fODF rescaling

The effectiveness of fODF rescaling in this study suggests that SNR measurements can be informative for resolving complex fiber compositions. The SNR measurements are likely region-sensitive because different WM regions have different complications in their microstructure. In this work we faced the complication that A-P oriented thick bundles bias the SNR in a frontal WM region, while in some other regions such as the corpus callosum this would not be the problem. A natural extension of the current study is to survey more WM regions and identify region-specific problems. For example, a voxelwise SNR map of the entire WM can be informative of potential fODF bias in different regions. Moreover, the method to estimate SNR can be further developed. We used cross-subject variability as an SNR indicator by taking advantage of the large sample size of the HCP data. Potentially the fODF variability among neighboring voxels can achieve similar goals if there is a fiber passing each voxel in the same direction. Given this reasoning the registration error across subjects is not a fundamental concern because nearby voxels contain similar information of the same underlying fiber.

### 4.4 Cross-species analysis for improving human tractography

NHP tractography plays a critical role in linking human *in vivo* neuroanatomy with the NHP *ex vivo* ground truth from the literature. On one hand, NHP tractography results can be directly compared with tract-tracing findings. Such comparison validates the efficacy of dMRI tractography methods in the same species. On the other hand, NHP tractography can be compared with human tractography to provide additional information that is otherwise inaccessible in human *in vivo* studies. In previous work, this strategy has been applied to elucidate topographical rules, fingerprints or network organization of anatomical connectivity in the human (Jbabdi et al., 2013; Neubert et al., 2015; Safadi et al., 2018; Tang et al., 2019). Here we further demonstrate that this cross-species, multimodal approach is effective in delineating detailed trajectories of individual bundles.

## Acknowledgement

J. G., S. F. and E. G. were supported by the National Institute of Biomedical Imaging And Bioengineering (NIBIB) of the National Institutes of Health (NIH) under Award Number R01EB027585.

## Supplemental Figures

**Supplemental figure 1.**
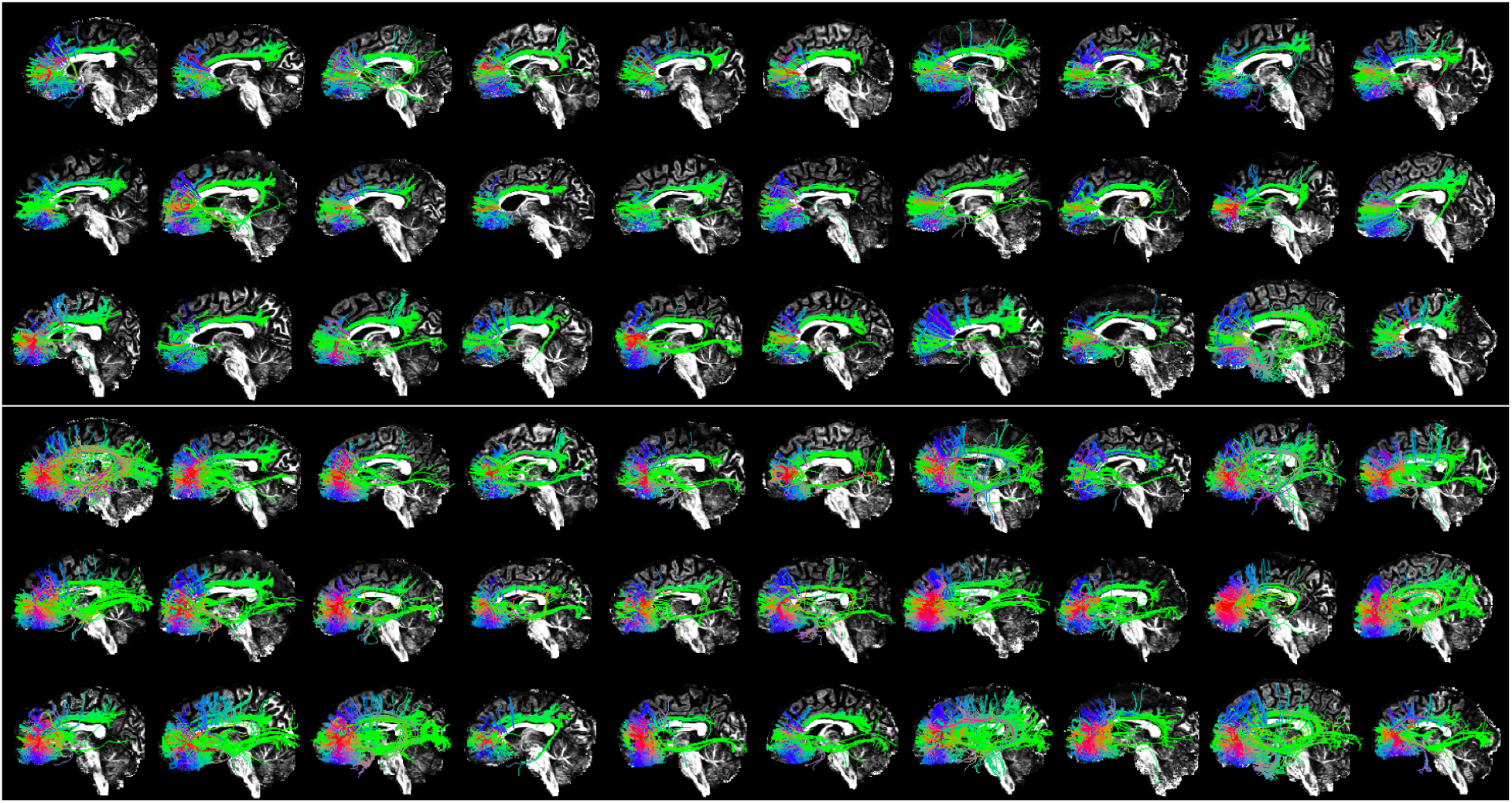
Tractograms of HCP subjects # 1-30. White line separates the results before (above line) and after (below line) fODF rescaling, with identical subject order.

**Supplemental figure 2.**
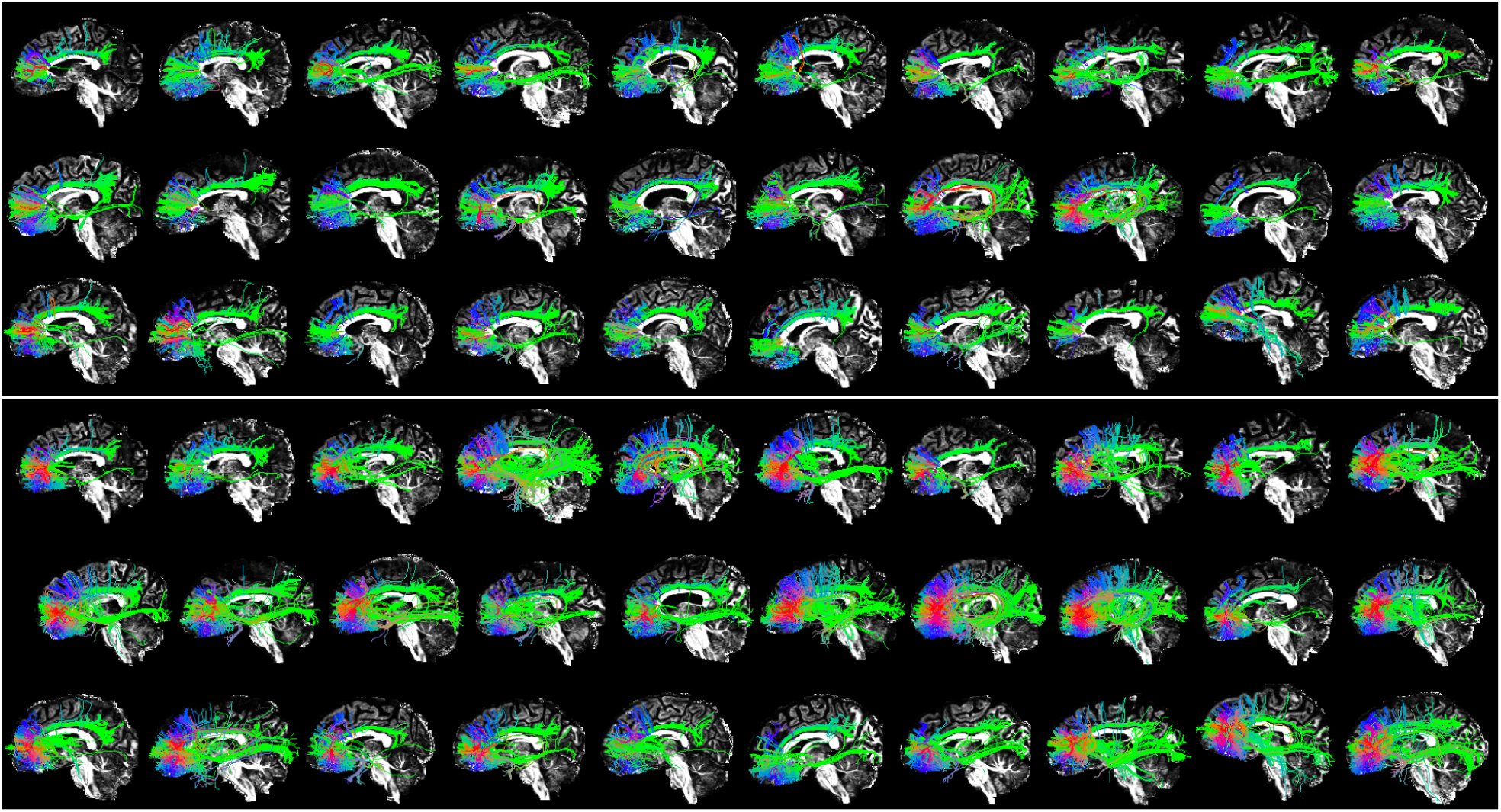
Tractograms of HCP subjects # 31-60. White line separates the results before (above line) and after (below line) fODF rescaling, with identical subject order.

**Supplemental figure 3.**
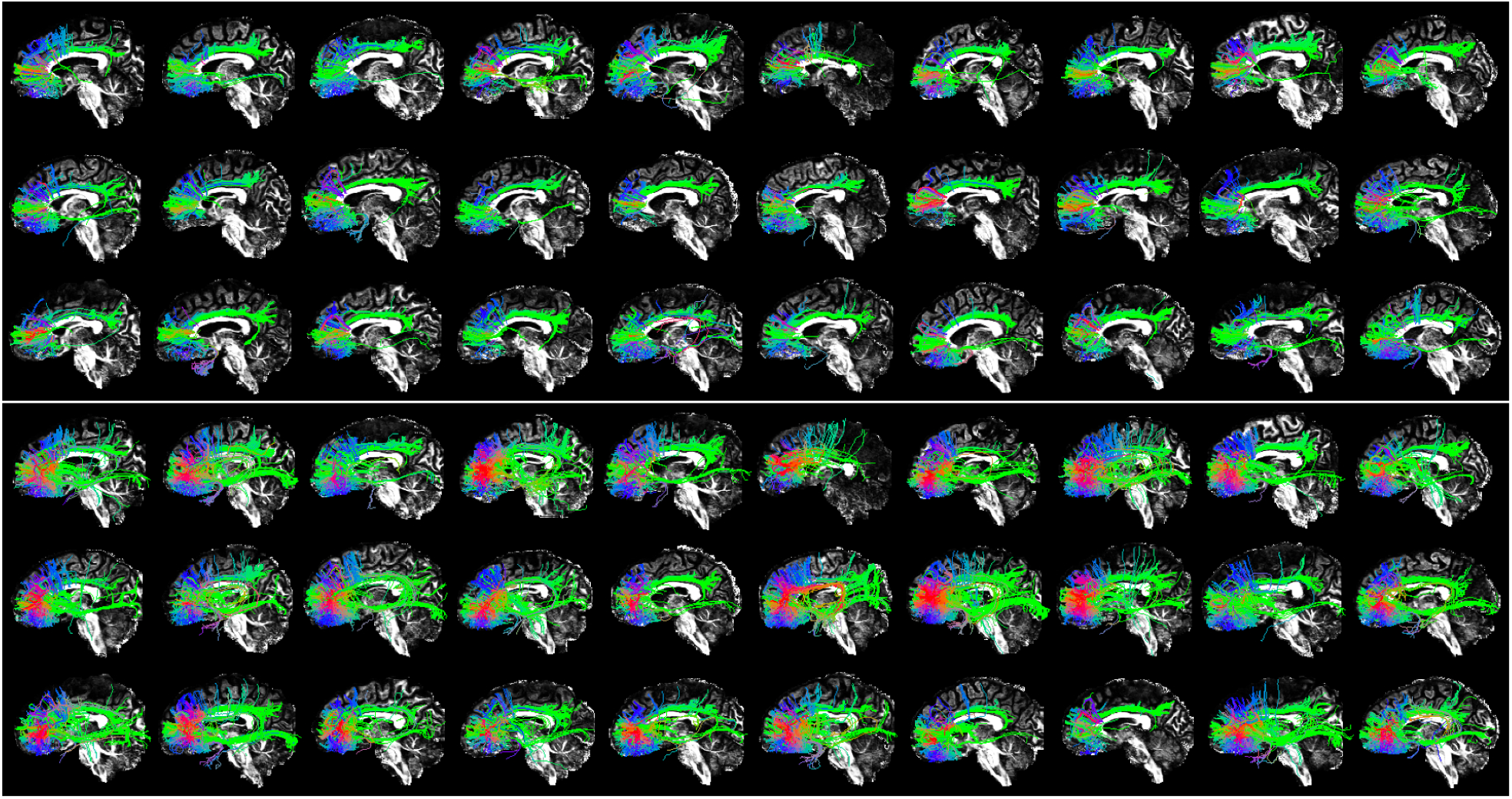
Tractograms of HCP subjects # 61-90. White line separates the results before (above line) and after (below line) fODF rescaling, with identical subject order.

**Supplemental figure 4.**
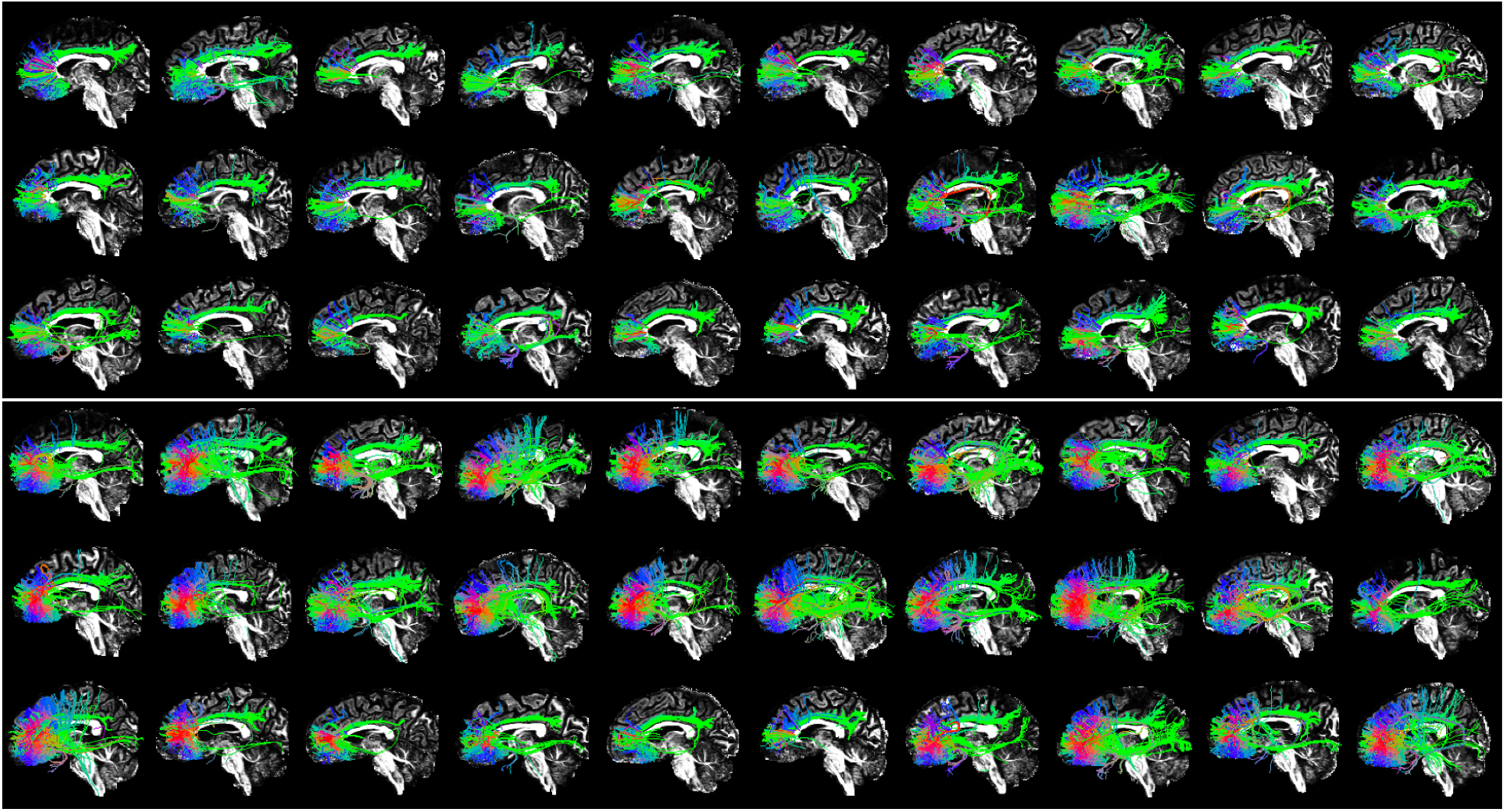
Tractograms of HCP subjects # 91-120. White line separates the results before (above line) and after (below line) fODF rescaling, with identical subject order.

**Supplemental figure 5.**
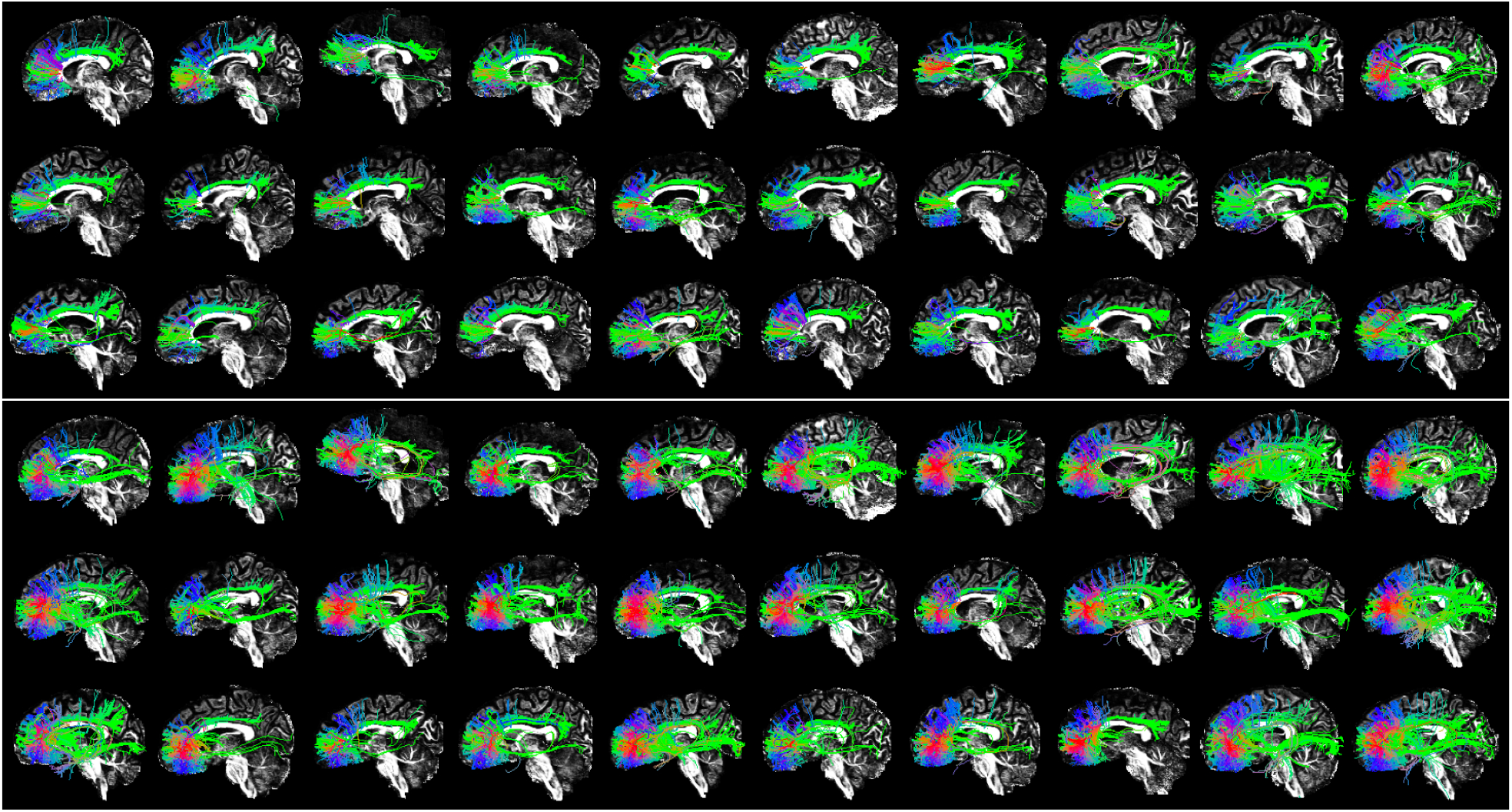
Tractograms of HCP subjects # 121-150. White line separates the results before (above line) and after (below line) fODF rescaling, with identical subject order.

**Supplemental figure 6.**
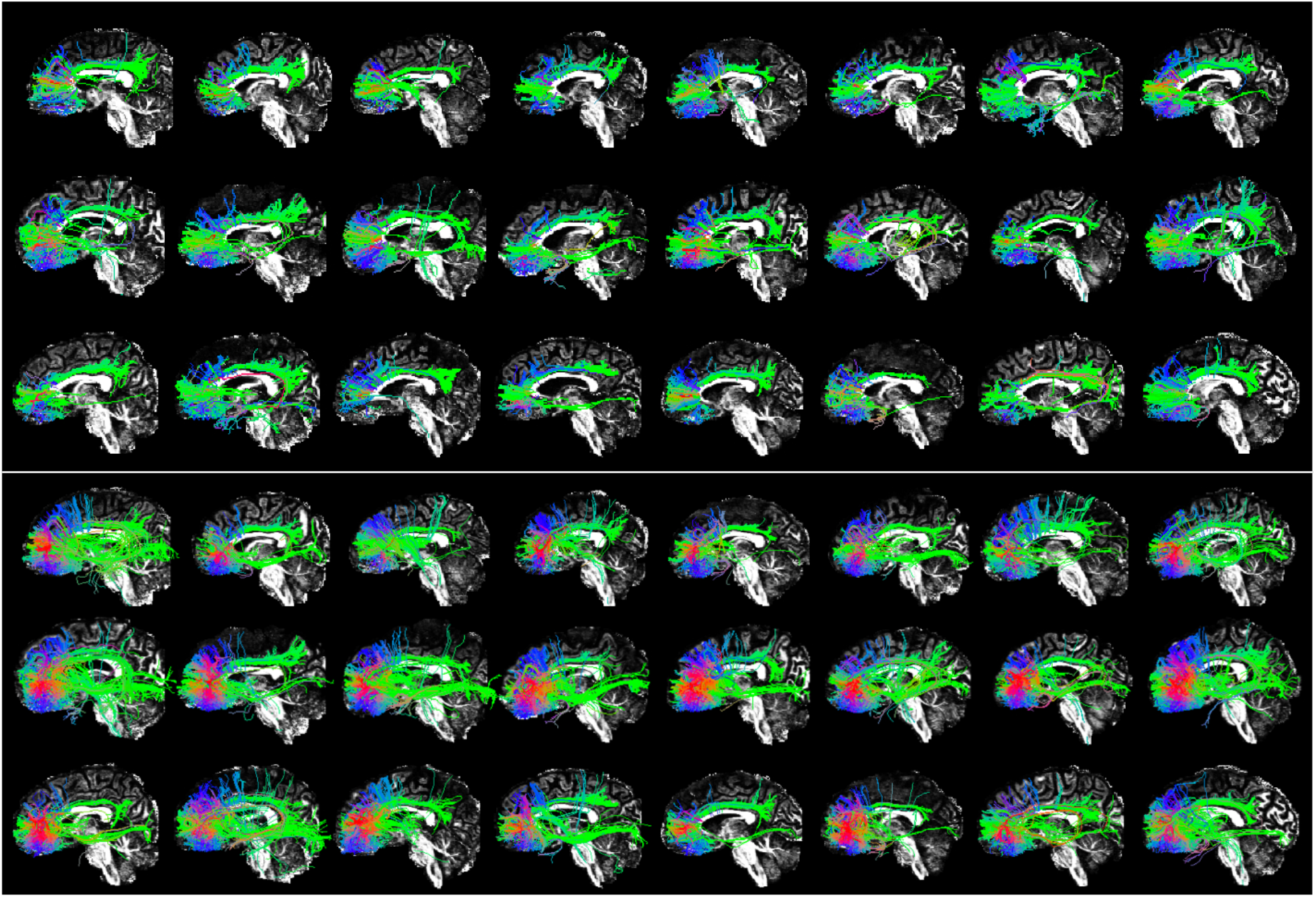
Tractograms of HCP subjects # 151-174. White line separates the results before (above line) and after (below line) fODF rescaling, with identical subject order.

## Notes

### Competing Interest Statement

The authors have declared no competing interest.

